# Electrotaxis disrupts patterns of cell-cell interactions of human corneal epithelial cells *in vitro*

**DOI:** 10.1101/2024.10.18.619085

**Authors:** Rebecca M. Crossley, Simon F. Martina-Perez

## Abstract

Electrotaxis, the process by which eukaryotic cells establish a polarity and move directionally along an electric field, is a well-studied mechanism to steer the migration of cells *in vitro* and *in vivo*. While the influence of an electric field on single cells in culture is well-documented, the influence of the electric field on cell-cell interactions has not been well studied. In this work, we quantify the length, duration and number of cell-cell interactions during electrotaxis of human corneal epithelial cells and compare the properties of these interactions with those arising in the absence of an electric field. We find that contact inhibition of locomotion and velocity alignment, two key behaviours observed during dynamic physical interactions between cells *in vitro*, are strongly affected by an electric field. Furthermore, we establish a link between the location of a cell-cell contact on the cell surface and the resulting cell interaction behaviours. By mapping the regions of the cell surface with a characteristic response to contact with another cell, we find that the spatial distribution of possible responses upon cell-cell contact is altered upon induction of an electric field. Altogether, this work shows how the electric field not only influences individual cell motility and directionality, but also affects cell-cell interactions.

## Introduction

Eukaryotic cells *in vitro* display intricate patterns of individual motion. At low densities, when cell-cell interactions are infrequent, single-cell motility is characterised as a stochastic process that takes into account random noise in the direction of travel, as well as persistence of motion as a result of subcellular signalling and cell morphology processes [1, 2]. External cues, such as electric fields, can be used to influence the characteristics of a cell’s individual motion [3–5]. The process of altering individual cell motility can provide valuable insights into how single cells aggregate information from their extracellular and intracellular environments to change their motility. Indeed, many eukaryotic cell types, such as human corneal epithelial cells, are known to exhibit electrotactic behaviour – targeted motion in the direction of an electric field. The ease with which electric fields can be used to steer cells has led to the acceptance of electrotaxis as a robust method of directing cell migration [6].

For example, previous work quantifying the impact of electric fields on single-cell motility demonstrated that the biased movement observed during electrotaxis was a results of polarity bias, where cells align their polarization direction with the surrounding electric field [7]. This study also used mathematical modelling to quantify the effect of this bias on the electrotactic phenotype. While the migratory properties of single cells have been previously studied and provide useful insights into the possible mechanisms driving their motility, cells in culture do not move in isolation. Even at cell densities as low as 100 cells per cm^2^, cell-cell collisions are frequent [7]. How cells interact with each other in such cases has been well-studied in the literature [8–10]. During cell-cell collisions of eukaryotic cells *in vitro*, various phenomena, such as contact inhibition of locomotion (CIL) [9, 10], and velocity of alignment of cells have been reported. These two phenomena are widely thought to play an important role in multicellular systems, such as the migration of cells in the neural crest [11], and in cancer [10]. Importantly, these behaviours vary between different cell types [9], suggesting that cells have a typical manner in which they respond to a cell-cell collision during migration.

The observation that cells exhibit characteristic responses to contacts with other cells, together with the finding that individual cell motility can be steered by applying an external electric field, invites the question of whether the application of an external influence, such as an electric field also alters the default pattern of cell-cell interactions. Previous studies of single-cell behaviours during electrotaxis have focused primarily on how individual cell motility is affected by the electric field [3, 4, 7], but a quantitative study of cell-cell interactions is still absent from the literature. In this study, we aim to answer the question of how the application of an external electric field influences cell-cell interactions.

To address these outstanding questions, we present an approach which quantifies and analyses cell-cell interactions from experimental brightfield images of human corneal epithelial cells from a previously published study [7]. Using automated segmentation, we carefully extract the locations of cells and their corresponding cell-cell interactions on the cell surfaces. We then use mathematical modelling techniques to identify distinct regions on the cell membrane that result in different behaviours after a cell contact, during both control conditions and when an external electric field is applied. Throughout, we demonstrate that the electric field not only influences the individual motility properties of human corneal epithelial cells, but also that it has a significant impact on the nature of cell-cell interactions *in vitro*. More generally, our methodology serves as a starting point to quantify cell-cell interactions in multicellular systems, and shows that there are fundamental differences in the patterns of cell-cell interactions in various extracellular conditions.

## Methods

### Electrotaxis experiment

Experimental data were collected by Prescott *et al*. [7] and consist of time lapsed videos of sparsely plated cells under two different experimental conditions, which we call the control and electrotaxis experiments. A detailed summary of the experimental procedures, methodology, materials and findings can be found in the original publication [7]. Initially, the telomerase-immortalised human corneal epithelial cells (supplied by Dr James Jester (University of California, Irvine)) were cultured to approximately 70% confluence at 37°C with 5% CO2 in EpiLife medium containing Ca2+ (60 *µ*M), which was supplemented with EpiLife defined growth supplement and 1%(v/v) penicillin/streptomycin (all materials purchased from Thermo Fisher Scientific (Waltham, MA)). In order to facilitate cell attachment for the subsequent experiments, cells between passages 55 and 65 were seeded at low density (100 cells cm^−2^) and cultured overnight (12 - 18 hours). During electrotaxis, an electric current was applied to the petri-dish chambers via an agar-salt bridge (agar purchased from MilliporeSigma (Burlington, MA)), which was connected using silver-silver chloride electrodes in Steinberg’s solution (58 mM NaCl and 0.67 mM KCl and 0.44 mM Ca(NO3)2, 1.3 mM MgSO4, and 4.6 mM Tris base (pH 7.4)). To ensure a strong salt bridge contact was maintained throughout the experiment, fresh cell culture medium (EpiLife) was regularly added into the reservoirs, where measuring electrodes at the end of the electrotaxis chamber connected the multimeter used to monitor the electric field strength.

During the subsequent experiments, cell migration was monitored using phase-contrast microscopy, namely with an inverted microscope (Carl Zeiss, Oberkochen, Germany) equipped with a motorized stage and a regular 10× objective lens. Time-lapse phase images were obtained at 5 minute intervals using Metamorph NX imaging software (Molecular Device, Sunnyvale, CA) throughout the 5 hour experiments. Under control conditions, no electric field was applied.

However in the electrotaxis experiments, cells were subjected to a 200mV mm^−1^ electric field from left to right between times 1 hour and 3 hours, which was then reversed and applied from right to left from 3 hours to 5 hours.

### Data

The dataset analysed in this work is derived from the before-mentioned experiment, which is described in [7] and investigates the impact of an electric field on cell motility. The original images from the associated experiments, which were used as raw data for these analyses, are publicly available online at https://zenodo.org/records/4749429.

### Cell segmentation and tracking

Using the raw image data from the original experiment [7], cells were automatically segmented using the pre-trained `cyto2` model with an object detection diameter of 25*µ*m in Cellpose [12]. Frame-by-frame visual comparisons between computed cell segmentations and experimental images revealed that no further manual correction of segmented images was required. Cell tracks and standard Cellpose fit features, such as best ellipsoid fit, aspect ratio, and cell orientation angle were obtained using the Fiji `Trackmate` plugin [13, 14] with the inbuilt Cellpose detector alongside the aforementioned settings, a LAP tracker with a max linking distance of 100*µ*m, and gap closing allowed over a maximum of 3 frames and 150 *µ*m. Visual inspection of the cell track outputs was used to inspect cell tracks shorter than the duration of the experiment, and were verified to correspond to cells leaving the field of view, or proliferation events. No further manual corrections were performed.

### Determination of cell-cell interactions

Given a segmentation mask containing labels corresponding to each of the different cells, we determined cell-cell contacts at every time point. Cell boundaries were determined as pixel locations directly adjacent to two different cell mask values, using the `skimage.graph` module as implemented in Python. Cells at further distances from each other were excluded as cell-cell contacts. All locations at which cell boundaries were directly adjacent were considered a cell-cell contact.

### Statistical tests

Statistical tests were conducted using an unpaired 2-sided Mann–Whitney nonparametric U test implemented in the `scipy` package in Python [15]. For time series, unless indicated otherwise, differences were computed point-wise in time.

## Results

### Electrotaxis directs single-cell motion along the electric field

As a first step towards understanding how the electric field influences the behaviour of human corneal epithelial cells, we quantified how single-cell migratory behaviours were affected after the electric field was turned on. The plots in the top row of Figure 1 demonstrate that the electric field re-directs motion in the direction parallel with the field. Cells in the control experiment, and cells in the electrotaxis experiment before the electric field is switched on (*i*.*e*. before *t* = 1h) move randomly, as shown by an average velocity in the *x* direction of zero, and an average directionality of zero. Here, the directionality, Φ, is defined as

**Figure 1.**
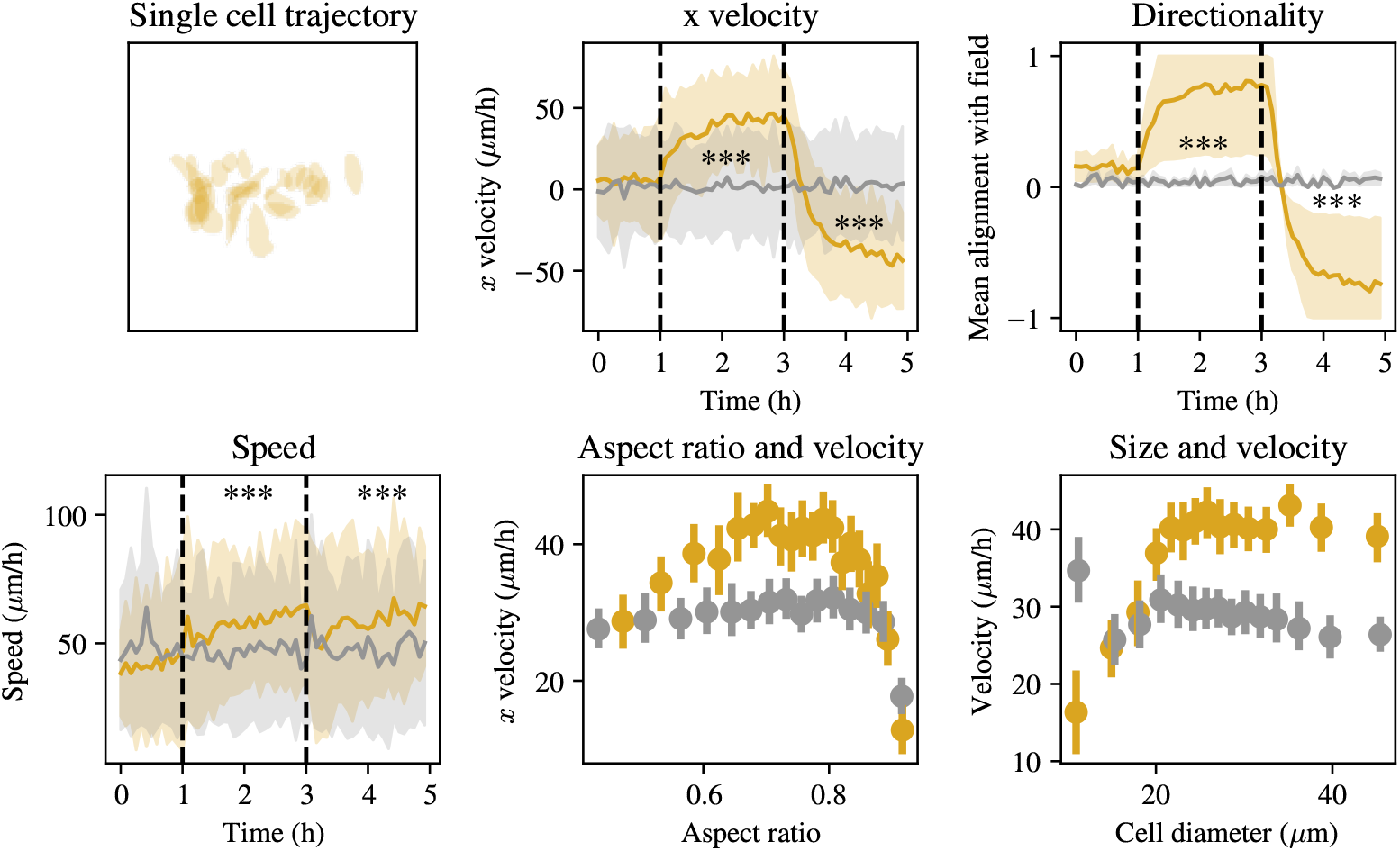
Top row, from left to right: plot of a individual cell’s trajectory during the electrotaxis experiment between *t* =0h and *t* = 1.5h; plot of the *x* component of the cell velocities against time, averaged over all cells in the control experiment (grey) and in the electrotaxis experiment (yellow), with 50 percentile results shaded; plot of the mean alignment of the cell with the electric field against time. Bottom row, from left to right: plot of the speed of cells against time; plot of the aspect ratio of a cell (*i*.*e*., the ratio of short axis to long axis on the modelled ellipsoid shape of the cell - see Methods) against the velocity of the cell; plot of the cell size against velocity of the cell. *** denotes a *p*-value less than or equal to 10^−3^.

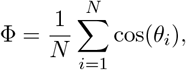

where *N* is the number of cells and *θ*_*i*_ is the angle between the velocity of cell *i* and the electric field. Once the electric field is applied, the directionality and the *x* component of the velocity of the cells in the electrotaxis experiment becomes positive. The directionality switches to negative in correspondence with the field reversing from right to left. Differences between electrotaxis and control experiment *x*-velocity and directionalities are significant at ≤ 10^−3^ for all time points at which the electric field is applied (see Figure 1 and Methods).

While the movement of the cells during electrotaxis is directed along the axis of the electric field, there is only a moderate but significant (at level 10^−3^ for all time points for which the field is applied, see Figure 1 and Methods) increase in speed of the cells. There is a marked overlap in one standard deviation from the mean between the speed curves in Figure 1, bottom row, left hand side. This indicates that, in this experiment, the electric field reorients the cells’ directionality without influencing their speed to the same extreme extent as seen in other (multicellular) systems at higher field strengths [16–18]. The fact that the cell speeds are of comparable magnitude in both experimental conditions then allows for a systematic comparison of their respective patterns of cell-cell interactions.

Finally, we ask whether any cell morphology characteristics influence the speed of singlecell migration velocity, both in control and electrotaxis conditions. By examining the aspect ratio of cells in comparison to their speed, we notice that there is an optimal aspect ratio of approximately 0.8 for cells to achieve maximum velocity during electrotaxis, whereas this effect is much less pronounced in control conditions. This observation is consistent with previous theoretical work relating cell shape to the efficiency of electrotaxis [19]. In both control and electrotaxis conditions, there is a noticeable drop in the speed of single cells as their aspect ratio approaches 1. This may be reflective of cells that are in the initial stages of mitosis: during prophase, cells slow down and stop migrating, whilst changes in the actin cytoskeleton occur that are often referred to as ‘rounding up’ [20]. During rounding up, cells collect their cell body contents into the center and produce a much more spherical shape, with a higher aspect ratio. This occurs whilst almost static, and could also explain the reduction in velocity observed at higher aspect ratios. Similarly, cell size influences migration velocity in control and electrotaxis conditions. As cell size increases, cell velocity increases, up to some maximum whereby increasing the cell size has minimal impact on the velocity. Interestingly, the control experiment appears to show small cells migrating at appreciably higher velocities than those during the electrotaxis experiment.

### Characterising cell-cell contacts during control and electrotaxis experiments

Having understood that the external electric field influences the motility behaviours of single cells, we wanted to understand whether the observed cell-cell contacts were affected during electrotaxis. To this end, we quantified key characteristics of cell-cell contacts. First, we analysed cell-cell contacts independently of cell motility, *e*.*g*., excluding information about the direction of cell motility. All statistics were computed in the same time interval 1h ≤ *t* ≤ 3h, which corresponds to the time period when the electric field is applied in the positive *x* direction during the electrotaxis experiment. For all comparisons, we used a Mann-Whitney U test to investigate the significance of the differences between the distributions.

Between control and electrotaxis conditions, the distribution of the lengths of cell-cell interfaces is unaffected (see Figure 2, difference between the length distributions is non-significant). The distribution of cell-cell contact duration additionally reveals that the vast majority of intercellular contacts are short-lived, though this distribution has long tails, indicating that some cells might stay in contact for prolonged periods of time, especially during electrotaxis (Figure 1, *p*-value of Mann-Whitney statistic is 2.20 × 10^−4^). No appreciable relationship exists between the contact length and the contact duration (data not shown). While the tails for cell-cell contacts are somewhat longer, the majority of cell-cell contacts occur on the same time scale during control and electrotaxis (see Figure 2). Finally, the size of migrating clusters is generally larger in electrotaxis than in control conditions (Figure 2, *p*-value for Mann-Whitney statistic 4.47 × 10^−5^). We reason that this might be due to cells undergoing electrotaxis moving in the same direction as one another, leading some contacts to be sustained due to an initial alignment in the direction of their velocities, whereas their individual motions would be stochastic and likely not aligned during control conditions. Together, these data show that the organisation of cell-cell contacts between human corneal epithelial cells is qualitatively similar in control and electrotaxis conditions, paving the way to investigate how such cell-cell contacts influence the way cells adjust their velocity upon an interaction with another cell.

**Figure 2.**
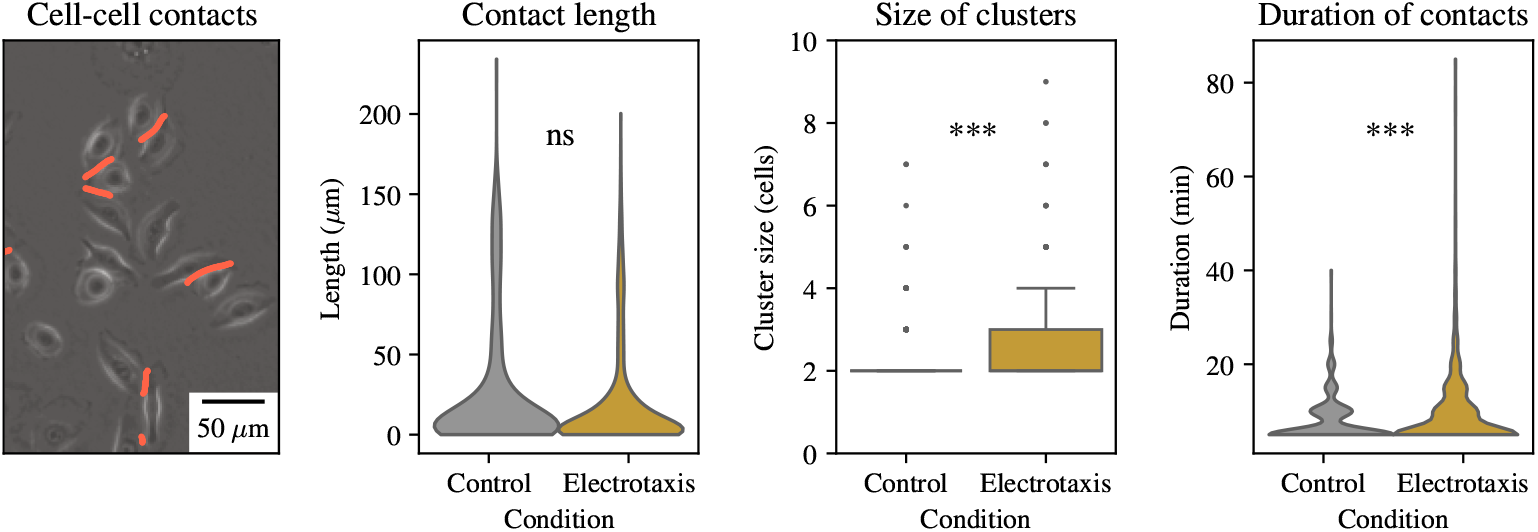
Cell clusters during unstimulated migration and electrotaxis. From left to right: phase image of human corneal epithelial cells at *t* = 1h of unstimulated migration with computed cell-cell contacts shown in red; distribution of contact lengths during unstimulated migration and electrotaxis (1h ≤ *t* ≤ 3h); number of cells in a cluster during unstimulated migration and electrotaxis (1h ≤ *t* ≤ 3h); distribution of the duration of cell-cell contacts during unstimulated migration and during electrotaxis (1h ≤ *t* ≤ 3h). ns denotes non-significance, *** denotes a *p*-value less than or equal to 10^−3^.

### Characterising cell-cell interactions

One key behaviour observed during cell-cell interactions across many different cell types [9–11] is CIL, which describes the reversal of a cell’s velocity to move directly away from contact with another cell. A number of different processes contribute to CIL including cell-cell recognition between specific cell surface molecules, signalling pathway initiation as a result of contact on the cell surface, and cytoskeleton rearrangement. The result of these processes generally is a change in the cell’s polarity and reversal away from the cell that it made contact with [10]. To quantify CIL between two cells given the computed cell tracks, we quantify the CIL of cell *i* with respect to cell *j* at time *t* as

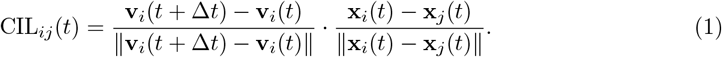

Here, **v**_*•*_, **x**_*•*_ are cell velocity and cell centroid location, respectively, and Δ*t* is the time increment between two successive frames in the dataset. The definition in Equation (1) emphasises how CIL depends on the component of the change in velocity of cell *i* in the direction of **x**_*i*_ − **x**_*j*_. Note here that CIL_*ij*_(*t*) ∈ [−1, 1], with a value of 1 corresponding to movement of cell *i* directly away from cell *j* and *vice versa* for a CIL of −1.

Another phenomenon arising in *in vitro* cell migration is that of velocity alignment. Here, alignment refers to the co-ordinated orientation of the cells making contact relative to one another. The interactions that we study in this work influence cell-cell alignment through direct physical contact with neighbouring cells. During physical contact, cell adhesion molecules such as integrins adhere junctions between cells to physically link them such that the cell cytoskeletons and matrices can align with one another. Alignment of cells can also occur as a result of interactions with environmental features such as adhesions with the extracellular matrix, mechanotransduction from nearby fluid flows, or other external signals such as growth factors or signalling molecules [1]. In this work, we quantify the degree of alignment arising from an interaction between two cells as the difference between their alignment pre-and post-contact, *i*.*e*.,

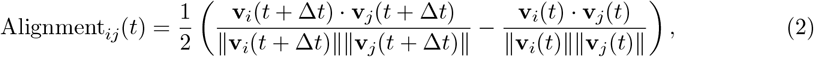

meaning that Alignment_*ij*_ ∈ [−1, 1], with a value of 1 indicating perfect alignment after a cell-cell contact, and a value of −1 indicating perfect anti-alignment. In what follows, we employ these metrics to understand the interactions between cells in control conditions, and how these interactions change in the presence of an electric field. Figure 3 shows a schematic of cartoon cell interactions corresponding to CIL and alignment.

**Figure 3.**
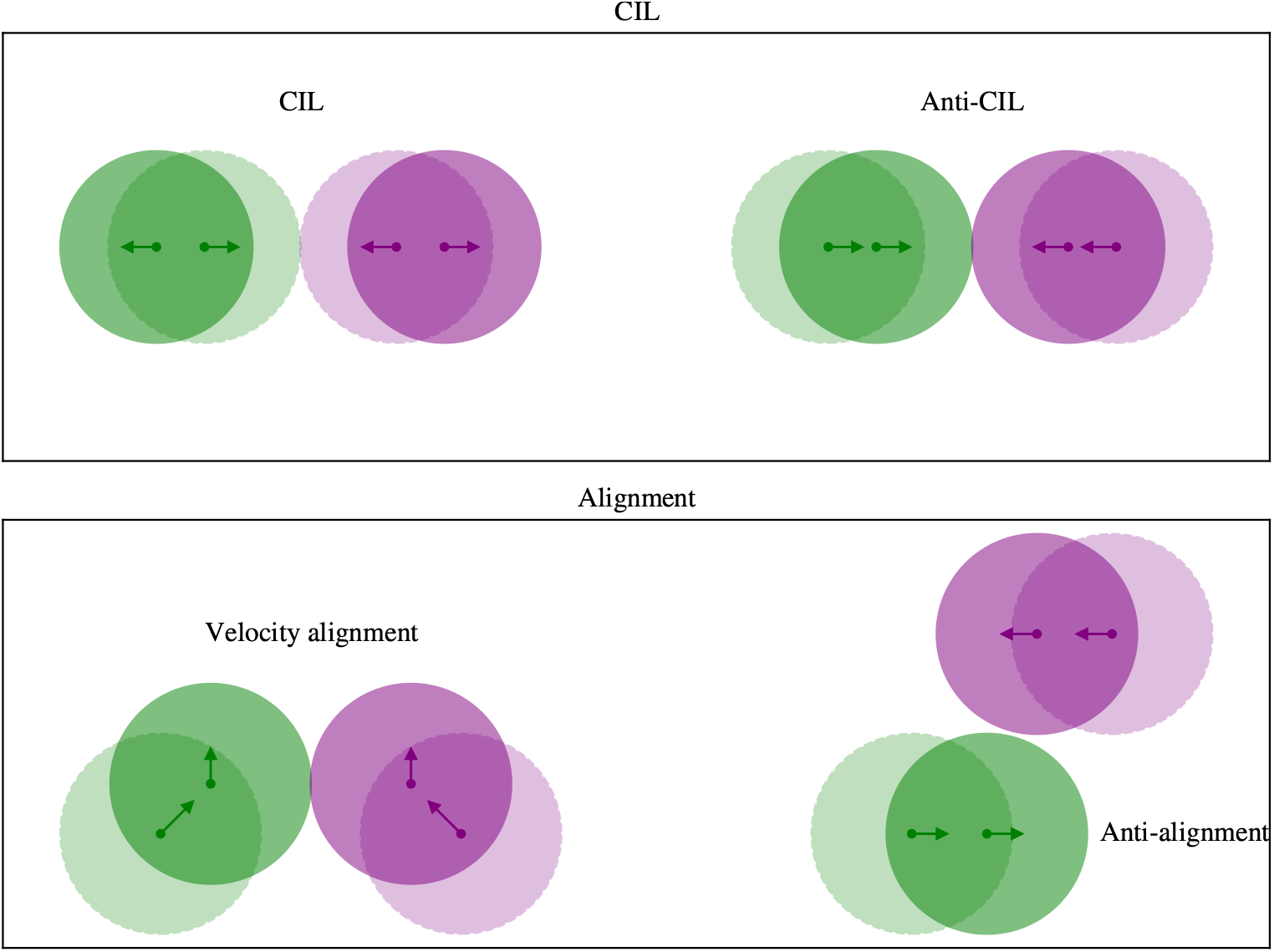
Cartoon cell interactions illustrating the mechanisms behind contact inhibition of locomotion (CIL) (top row) and alignment (bottom row). These metrics are defined in Equations (1) and (2), respectively. In all cartoons, two cells have an initial positions (hatched fill) which changes to a new position (solid fill) and the velocity over the next time step is indicated by an arrow. Top: left shows an interaction with CIL, and right shows an interaction with-out CIL (anti-CIL). Bottom: left shows an interaction with two cells aligning their velocities, corresponding to a positive value of the alignment metric in Equation (2), whereas the right interaction shows two cells that are perfectly anti-aligned.

### CIL and alignment during the control experiment

To establish patterns of cell-cell interactions in the absence of an electric field, we first sought to quantify the effect of cell-cell collisions on the subsequent movement of individual cells during unstimulated migration. We did this by quantifying CIL and relative alignment of interacting cells, as defined in Equations (1) and (2). Figure 4 shows that cell-cell collisions are frequent and show a wide range of locations along the cell surface where contacts take place. Inspired by this, we sought to develop a systematic way to quantify the *position* of a cell-cell contact on the cell surface of two interacting cells. Using an ellipse fitted to each cell (see Methods), we project each contact site on the cell surface to its closest site on the corresponding ellipse. We then identify the front of the cell as the short axis in the direction aligned with the velocity because human corneal epithelial cells become highly polarised and move in the direction of their short axis in both control and electrotaxis experiments. Subsequently, we compute the angle, *ϕ* ∈ [0, *π*], between the front of the cell and the computed site of contact (*i*.*e*. we obtain a left-right symmetric measure for the location of a contact). This allowed us to robustly quantify the location of cell-cell contacts across all interactions.

**Figure 4.**
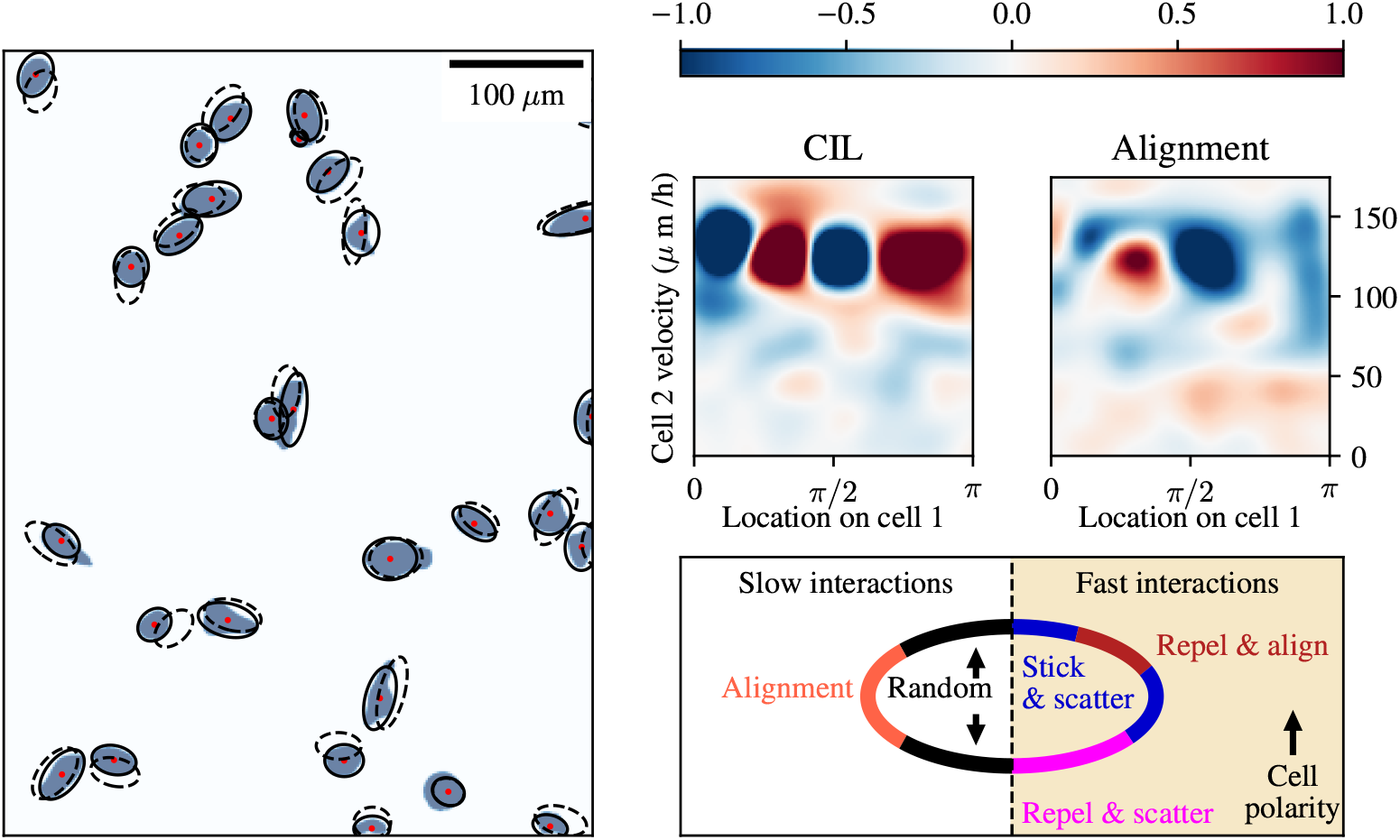
Dynamic interactions between cells in control conditions. Left: cell masks and fitted ellipses (solid lines) corresponding to the same frame as the segmentation masks. Ellipses with dashed lines correspond to fitted ellipses in the next frame. Top right: heat maps showing a measure of the contact inhibition of locomotion (CIL) and alignment against the location the contact was made on cell 1 and the speed of the contacting cell 2. Bottom right: schematic demonstrating the different types of interactions observed when contact occurs in different locations on the cells at slow and fast velocities.

First, we investigated the response of a cell – which we call the *focal cell* or cell 1 – as it makes contact with another cell – which we call the *neighbour cell*, or cell 2. Since cell-cell interactions are not necessarily reciprocal, each interaction will yield data for its cells playing the role of both focal cell and neighbour cell. Interactions with cells migrating at a low speed – defined here as lower than 50*µ*m/h for the neighbour cell – generally lead to no appreciable pattern in CIL or relative alignment (Figure 4, top row): regardless of spatial location both the CIL and relative alignment measures for the focal cell are close to zero. This finding suggests that a minimum speed is required for non-negligible interactions. The relative alignment heatmap in Figure 4 shows that there is a slight increase in alignment on average when the contact site is in the region around *ϕ* = *π/*2, *i*.*e*., to the side of the cell.

Interactions with a neighbour cell migrating at a high speed – defined here as higher than 100 *µ*m/h – result in a different pattern of interaction. By observing the distinct pattern in the heatmaps of Figure 4, we summarise these behaviours in four representative regions of the cell that correspond to different behaviours. In the region closest to the front of the cell, we find that, for sufficiently high migration speed of the interacting cell, no CIL is observed. This effect occurs regardless of the velocity of the focal cell. Rather, there is a negative value of the computed CIL metric, indicating that cells do not actively move away from the site of cell-cell contact, but continue moving in their original direction of travel. This result is indicative of the persistence of the focal cell’s motion, which ensures the cell continues moving directionally, even after cell-cell contact has been established. In this region, we also observe a change in alignment of the cell in the opposite direction to the migratory direction of the neighbour cell it has come into contact with. Here, we describe this behaviour as ‘stick & scatter’, as cells initially stick to one another but reorient away from one another after coming into contact. This same behaviour is also observed in the region corresponding to the side of the cell.

Interactions upon contact in the the region diagonal and to the front of the focal cell, lead to repulsion and alignment of the focal cell with respect to its neighbour (see schematic in Figure 4). These interactions are characterised by the focal cell quickly moving away from its neighbour (as indicated by the value of CIL close to 1), while its velocity aligns to that of the neighbouring cell. This pattern is opposite from that occurring toward the rear of the focal cell (‘repel & scatter’ in the schematic in Figure 4). This ‘repel and scatter’ pattern of interaction is physically intuitive: contacts in this region are only made when one of the interacting cells is moving faster than the other, and the interactions are front-to-back. Due to the persistence in the motion of the faster moving cell, no CIL is observed. At the same time, the interaction effectively pushes the front, focal, cell to align with the velocity of the neighbour cell with faster motion, leading to relative alignment.

In sum, cell-cell interactions during control conditions display a dependence on the velocity of the neighbouring cell as well as on the location of the interaction on the focal cell surface. Importantly, we identify different representative regions along the focal cell surface that correspond with different types of cell-cell interactions. This finding complements existing studies that suggest that the relative positions of cells in an interaction determine the effect of the interaction on collective motion. Previously, for example, LaChance *et al*. identified that cells in epithelial sheets only display correlated motion with a subset of neighbours which are to their front [21]. Further work also demonstrated that genetic perturbations within the same cell type can lead to dramatic alterations to these patterns of intracellular coordination [22].

### Interactions during the electrotaxis experiments

Having understood the patterns of cell-cell interactions during control conditions, we turn to the question of how the electric field impacts cell behaviours. As in control conditions, the type of cell-cell interaction varies dramatically depending on the speed of the interacting neighbour cell (Figure 5, top right). If the speed of the neighbour cell is low (below 50*µ*m/h), and contact occurs at the front or rear of the focal cell, we observe alignment of the focal cell toward the velocity of the neighbouring cell and no CIL. Contact at the side of the focal cell, in contrast, leads to CIL and relative alignment of the focal cell with respect to the velocity of the neighbour cell. During electrotaxis cells are exposed to a migratory command influencing their direction of travel.

**Figure 5.**
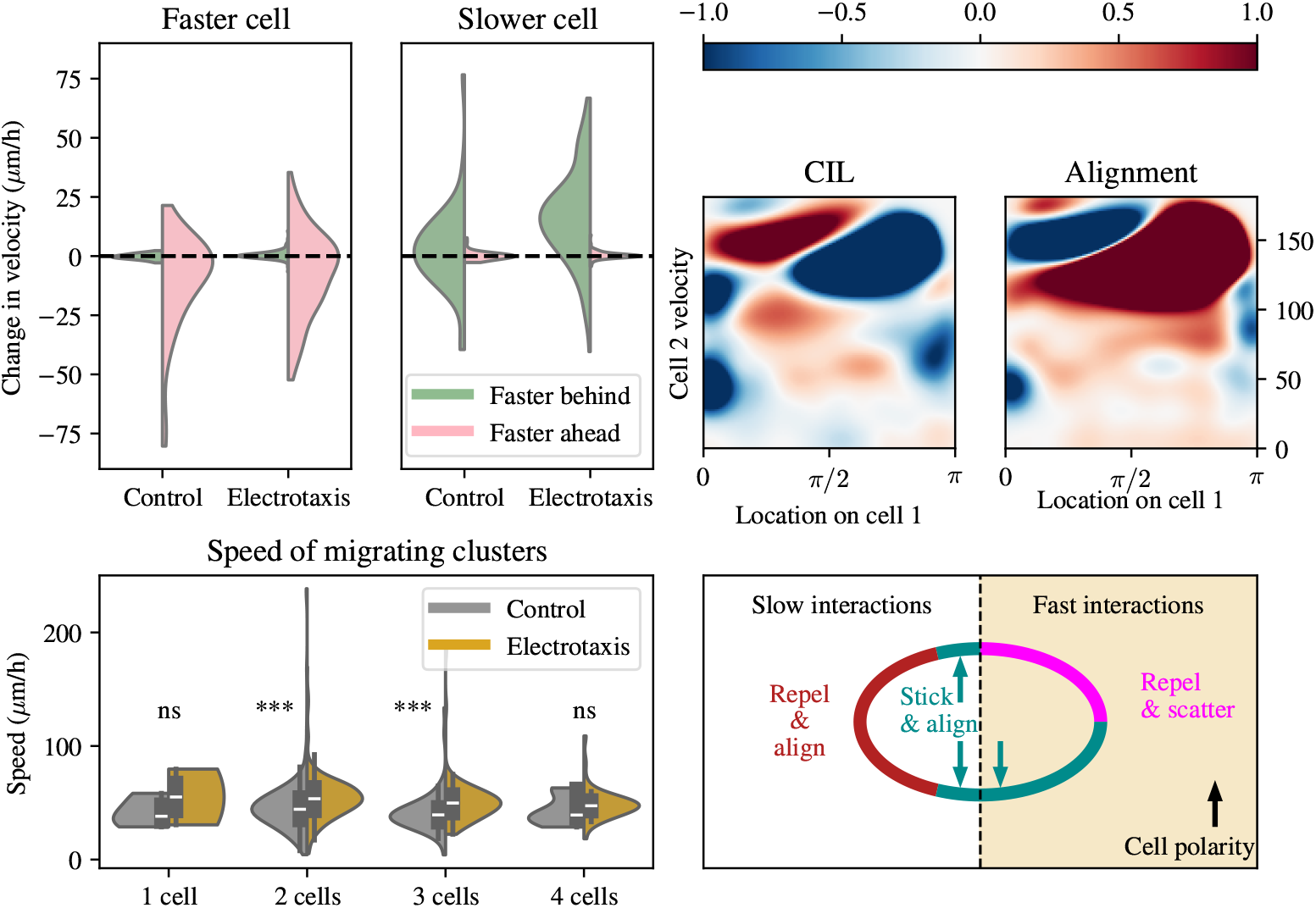
Dynamic interactions between cells during the electrotaxis experiment. Top left: violin plots showing distributions of velocities classified by the position (front/back) of the faster and slower moving cells. Top right: heatmaps showing a measure of the contact inhibition of locomotion (CIL) and alignment against the location the contact was made on cell 1 and the speed of the contacting cell 2. Bottom left: violin plots of the speed of migrating clusters of cells in control and electrotaxis experiments. Bottom right: schematic demonstrating the different types of interactions observed when contact occurs in different locations on the cells at slow and fast velocities. ns denotes non-significance, whereas *** denotes a *p*-value less than or equal to 10^−3^.

At higher speeds (above 100*µ*m/h), the heat maps in Figure 5 reveal the same ‘stick and align’ behaviour observed under control conditions when a contact occurs at the rear of the focal cell. In this case, the neighbour cell making contact in the rear is most likely travelling much faster than the front cell and therefore a positive value for the CIL metric defined in Equation (1) is observed: the cell is pushed forward. As a result, the focal cell at the front also aligns with the neighbour cell toward its back. It should be noted here that, while the focal cell indeed moves away from the contact with its neighbour, this does not constitute CIL in the traditional sense, as there is no repolarisation away from the cell-cell contact. Conversely, contacts occurring on the front of the focal cell cause repulsion and anti-alignment with respect to its neighbouring cell, (see the ‘repel & scatter’ region in the schematic of Figure 5). These collisions represent a faster moving focal cell making contact with the rear of a slower moving neighbour cell, leading to the faster cell slowing down. The effect of this is that there is no apparent CIL and relative anti-alignment.

To further understand the dynamics of front-back interactions during electrotaxis, we analyse the changes in velocity of the focal cell immediately post-collision whenever this collision is front-to-back (Figure 5, top left). If in such an interaction the faster moving cell is behind the slower moving cell, the velocity of the slower cell increases on average, as a result of the collision in electrotaxis, but not in control conditions (Figure 5). So, during electrotaxis, the slower cell at the front of the cell-cell contact is being pushed along and sped up by contact with a faster cell from behind, whose velocity is largely unchanged by the interaction. The variance in the velocity changes are markedly smaller than for the slower cell. If, instead, the faster cell is ahead when a contact occurs, its velocity remains unaffected on average, although the variance of this distribution is far greater. In contrast, the slower moving cell does not exhibit any appreciable change in velocity. This effect is equal in both control and electrotaxis. Thus, interactions with the slower cell behind can be understood as only the faster cell being slowed down, leading to overall velocity changes for the faster cell, but not for the slower cell.

Finally, to understand whether the effect of cells speeding and slowing each other depending on their relative positions leads to larger clusters of cells to move at different speeds than slower cells, we analysed the speed of migrating clusters of cells in both control conditions and electrotaxis (Figure 5, bottom left). The speed of migrating clusters does not change significantly for single cells or large clusters of four cells. It changes significantly for clusters of two or three cells (with *p*-values for the Mann-Whitney statistic given by 5.21 × 10^−11^ and 1.89 × 10^−5^, respectively), but this increase is modest, with the means for both groups being less than 15% apart. This suggests that, at least for small clusters of cells, clustering does not lead to overall coordination and efficiency of migration.

### Impact of the electric field reversal during electrotaxis experiments

Having established that the electric field creates a signal resulting in the rapid polarisation of the cells, we wondered how the instantaneous switch in the direction of the field at *t* = 3 hours would affect the movement of cells.

Previously, experiments in confluent cell sheets have shown that cells display U-turns upon field reversal [23]. To assess whether cells in this experiment turn by instant polarity reversal or by U-turning, we investigate the velocity in the direction perpendicular to the field, *i*.*e*., *v*_*y*_, in Figure 6. If there were U-turns on the order of greater than 5 minutes (the time interval between frames), we reason that there would be distinct patterns of cells displaying nonzero absolute values of *v*_*y*_ in a period post-reversal before returning to zero, on average. The data in Figure 6, however, does not show such a trend, and, in fact, show that the distribution of velocities is unchanged before and after field reversal (Figure 6, difference between distributions is non-significant). This is indicative of cells quickly reversing their polarity and migrating in the opposite direction, or making a U-turn faster than the time scale at which the data are acquired. We propose that at low densities, cells can reverse their polarity extremely fast, resulting in a near-instantaneous reversal of their velocity, whilst in highly confluent cell systems the physical coupling of cells could create more complicated spatial velocity patterns, such as topological defects [24].

**Figure 6.**
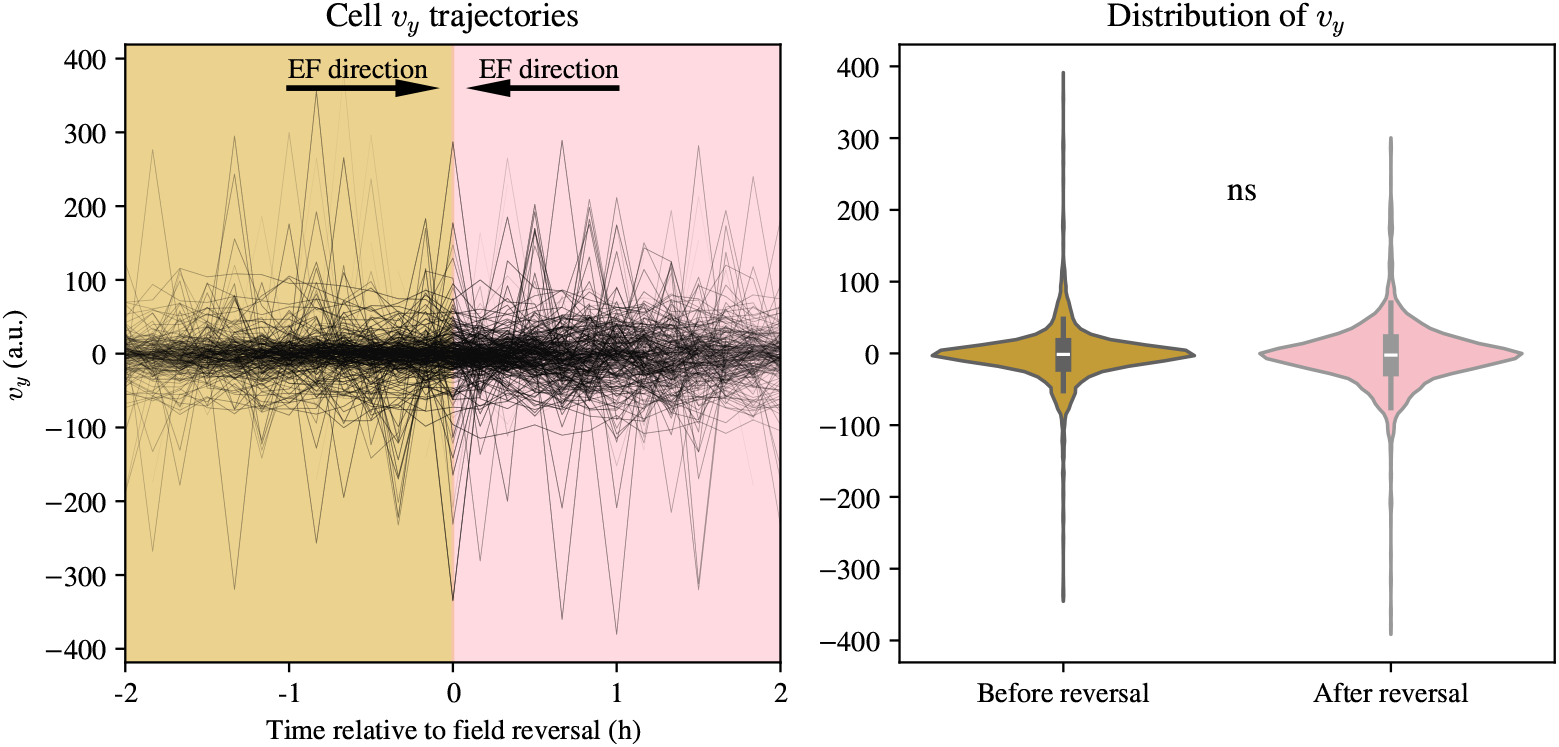
Velocities of cells in the direction perpendicular to the electric field (EF) before and after field reversal. Left: *v*_*y*_ for individual cell trajectories against time. Right: violin plots of the distribution of *v*_*y*_ pre- and post-field reversal. ns denotes non-significance.

We could postulate various potential mechanisms behind the rapid polarity shift observed when changing the direction of the electric field. Comprehensive reviews of the impact of electric fields on cell behaviours, such as [25], highlight the role of intracellular signalling pathways, or charges on cell-surface receptors as hypothetical factors determining the directionality of cell migration during electrotaxis. In particular, the actin cytoskeleton of a cell undergoes dynamic reorganisation during electrotaxis, with Rho GTPases such as RhoA and Cdc42 playing key roles in modulating contractility and actin polymerization, respectively [26]. Alternatively, signal transduction pathways may further amplify spatial cues from electric fields, enabling swift directional changes, which have also been modelled in the theoretical framework of directional sensing [27].

## Discussion

By quantifying the changes in individual cell motility after cell-cell collisions, this work addressed the question of how an external electric field influences the patterns of cell-cell interactions between migrating human corneal epithelial cells. While the average length and duration of cell-cell contacts were hardly influenced by the electric field, there was a pronounced difference in how cells interacted with their neighbours when the field was turned on. Both in control and electrotaxis conditions, how cells re-polarise and align with the cells they physically interact with, depends on the physical location on the cell surface at which contact is made, as well as the velocity of the interacting neighbour cell. These locations essentially create a partition of the cell surface in different zones such that contact at these zones results in characteristic motile responses, as demonstrated in Figure 7. Strikingly, turning on an external electric field changes both the spatial location and the nature of these zones, indicating that the electric field might not only influence single-cell motility, but also the transduction of external mechanical factors relating to a cell-cell collision.

**Figure 7.**
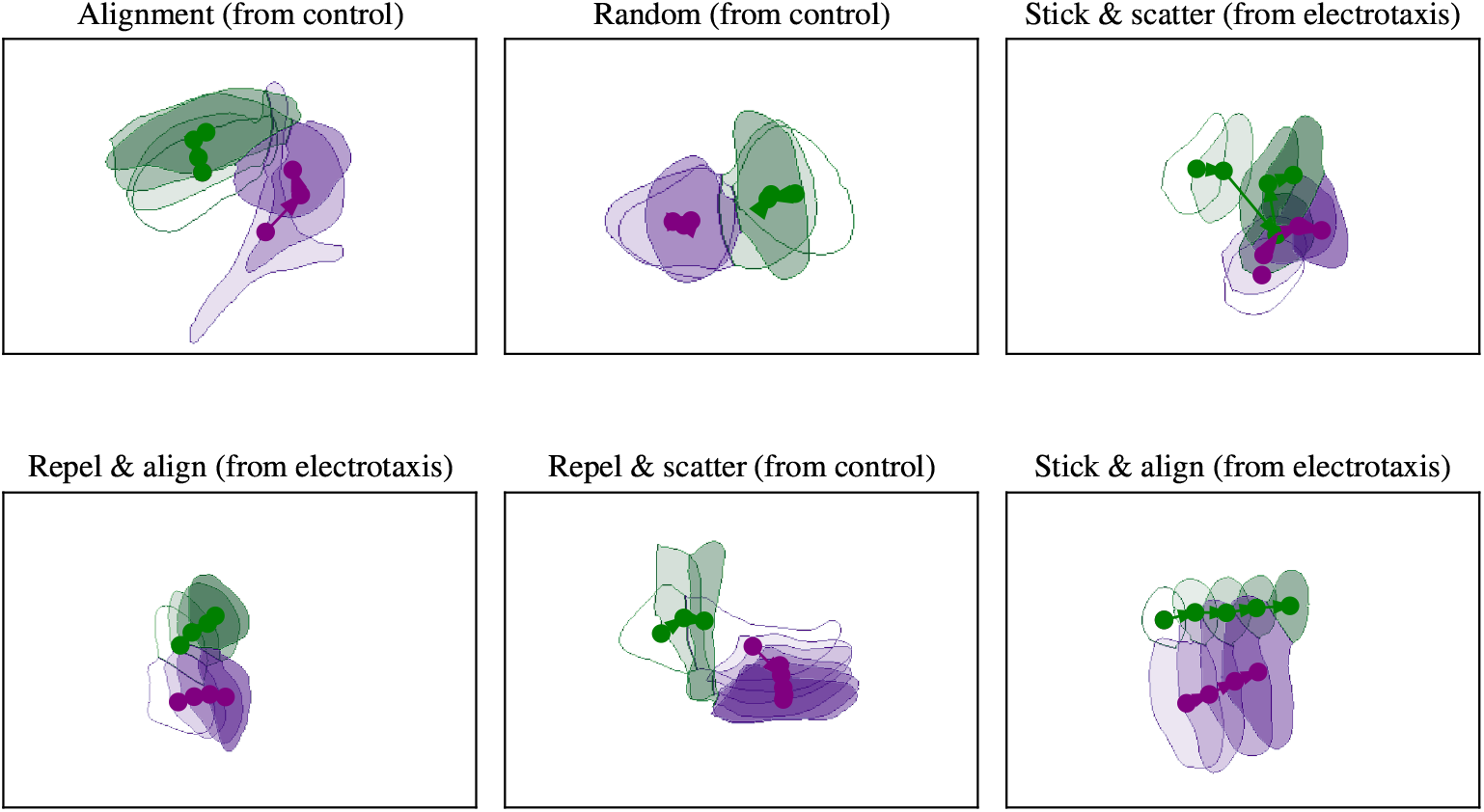
Cell masks of some representative interactions between two cells, labeled according to the schematics in Figures 4 and 5. Note that, as shown by the schematics in Figures 4 and 5, some behaviours are characteristic of the control or electrotaxis experiments, respectively. See also Table 1 for a definition of the names of the interaction types shown in this figure.

With these findings, we extend the existing literature which primarily investigates single cell behaviours during electrotaxis, and guided cell migration more generally. Despite the question of how an electric field influences the motility of single cells and cellular collectives having received ample attention in the literature, the question of how the electric field imposes a signal that overrides the intrinsic patterns of cell-cell interaction has not yet been considered. This study provides a starting point to suggest that an external cue can override normal cell responses to contact with other cells, which goes beyond exclusively guiding cells in the direction of the field, thus raising several interesting directions for follow-up studies.

**Table 1:** Glossary of important terms.

| Term | Definition |
| --- | --- |
| Alignment | The co-ordinated orientation of cells towards one another after making contact. |
| Contact inhibition of locomotion (CIL) | The reversal of a cell's velocity to move directly away from contact with another cell. |
| Stick & scatter | After a cell-cell interaction, cells do not actively move away from one another (CIL), but do change alignment to move away from the migratory direction of the cell it has come into contact with. |
| Stick & align | CIL and alignment of cell velocities are observed after a cell-cell interaction. |
| Repel & scatter | Post cell-cell contact, cells move away from one another in opposing directions (no CIL, no alignment). |
| Repel & align | No CIL is observed after a cell-cell contact, but alignment of the cells velocities occurs. |

The finding that patterns of cell-cell interactions are altered upon the imposition of an external electric field invites further study into the mechanistic origins of such cell-cell interaction patterns. For example, while the signalling networks that lead to single-cell migration in the presence of an electric field are somewhat understood and known to be shared with those implicated in chemotaxis [28], a straightforward question arising is whether the signalling networks known to be involved in *e*.*g*., CIL are impacted by the electric field. Another avenue of investigation to explore why there is a marked change in cell-cell interactions would be to quantify any changes in sub-cellular organisation in response to an external electric field. For example, is there a difference in actin polymerisation or cytoskeletal organisation that could explain any mechanical differences leading to altered patterns of cell-cell interactions?

In this work, such questions cannot be interrogated since brightfield imaging at low magnification does not allow for detailed mapping of the cellular projections extended by the cells, nor of any sub-cellular contents. However, more advanced imaging modalities, including multichannel imaging to include actin and other structures, or multi-omics approaches to unpick cells transcriptional and metabolic activities, could provide a sensible extension to unpick these questions. The answers to these questions form a starting point for enquiry into the causes of altered cell-cell interaction patterns.

Of relevance to this particular study is the fact that the mechanical properties of many cell types are affected by a wide range of environmental factors. For instance, in culture, the morphology and motility of most cell types are also known to depend significantly on the stiffness of the substrate they are plated upon [29–33]. Consequently, it would be valuable to investigate whether the composition of the medium used in these experiments influences the observed results and to explore how variations in substrate stiffness might affect the mechanisms or strength of directionality observed during electrotaxis. Introducing such key variables that determine cellular mechanical behaviour can be used in future studies into cell-cell interactions in electrotaxis to better tell apart the role of the electric field in influencing cell-cell interactions, and understand how it might interplay with other environmental cues. Doing this would increase our understanding of the optimal experimental conditions for this cell type, possibly advancing the application of electrotaxis in areas such as regenerative medicine, tissue engineering, and disease treatment.

The methods in this work can readily be extended to the analysis of cell-cell interactions in different cell types, and doing so can help establish a more systematic connection between the external electric field and its interference with standard cell-cell interaction patterns. Furthermore, there are several possible extensions to this work that one can consider. Firstly, this work only considers pairwise interactions between cells. While this type of interaction represents the vast majority of cell-cell interactions observed in this experiment, large clusters of cells are also regularly observed in multicellular systems seeded at higher density. Secondly, the effect of the electric field on the proliferation of eukaryotic cells merits direct attention. The phenomenon of ‘go-or-grow’ [34], in which there is an apparent energetic trade-off between proliferation and motility, has been studied and debated in several model systems [35–38]. Given that proliferation and electrotaxis are both energetically *expensive* events, understanding to what extent electrotaxis influences the proliferative properties of different cell types might inform our understanding of the ‘go-or-grow’ hypothesis, and how the electric field influences the control behaviours of eukaryotic cells, more generally. For a quantification of such effects, one would need data collected on sufficiently long time scales to observe the effects of the field on motility and proliferation: we observed in this experiment that the average time the cells spend in the mitotic phase to be in the order of 30 minutes, meaning that not enough data is available when electrotaxis is only performed for two hours on low density seeded cells. For this reason, the analysis of proliferation events necessitates the collection of electrotaxis data for a longer duration, which in turn might allow the quantification of other cell-cell interaction processes over longer time scales.

## Authors’ contributions

R.M.C and S.M.-P. conceived the project. S.M.-P. wrote the computational scripts used for data analysis. R.M.C. and S.M.-P. analysed the data and interpreted the results. R.M.C. and S.M.-P. wrote the article. The authors contributed equally and their names appear in alphabetical order. Both authors agree that either may list themselves first on a CV.

## Declaration of competing interest

The authors do not have any competing interests to declare.

## Acknowledgements

The authors would like to thank Prof. Ruth Baker for her useful insights and discussions regarding the manuscript. S.M.P. would like to thank the Foulkes Foundation for funding. R.M.C. would like to thank the Engineering and Physical Sciences Research Council (EP/T517811/1) and the Oxford-Wolfson-Marriott scholarship at Wolfson College, University of Oxford for funding.

## Notes

https://zenodo.org/records/4749429

### Competing Interest Statement

The authors have declared no competing interest.

### Summary of Updates

Added schematics and some minor additions to text.

